# A G-protein-biased S1P_1_ agonist, SAR247799, improved LVH and diastolic function in a rat model of metabolic syndrome

**DOI:** 10.1101/2021.09.14.460397

**Authors:** Maria-Francesca Evaristi, Bruno Poirier, Xavier Chénedé, Anne-Marie Lefebvre, Alain Roccon, Florence Gillot, Sandra Beeské, Alain Corbier, Marie-Pierre Pruniaux-Harnist, Philip Janiak, Ashfaq A. Parkar

**Affiliations:** Sanofi R&D, Diabetes and Cardiovascular Research, 1 Avenue Pierre Brossolette 91385 Chilly-Mazarin, France; Sanofi R&D, Molecular Histology and Bioimaging Translational Sciences, 1 Avenue Pierre Brossolette 91385 Chilly-Mazarin, France; Sanofi R&D, Biomarkers and Clinical Bioanalyses, Translational Medicine and Early Development, 371 Rue Professeur Blayac 34184 Montpellier, France; Sanofi US Services, Diabetes and Cardiovascular Research, 55 Corporate Drive, Bridgewater, NJ 08807, USA

**Keywords:** endothelium, HFpEF, SAR247799, S1P_1_

## Abstract

**Aim:** Heart failure with preserved ejection fraction (HFpEF) is a major cause of death worldwide with no approved treatment. Left ventricular hypertrophy (LVH) and diastolic dysfunction represent the structural and functional components of HFpEF, respectively. Endothelial dysfunction is prevalent in HFpEF and predicts cardiovascular events. We investigated if SAR247799, a G-protein-biased sphingosine-1-phosphate receptor 1 (S1P_1_) agonist with endothelial-protective properties, could improve cardiac and renal functions in a rat model of metabolic syndrome LVH and diastolic function.

**Methods:** 31- and 65-week-old obese ZSF1 (Ob-ZSF1) rats, representing young and old animals with LVH and diastolic dysfunction, were randomized to a chow diet containing 0.025% (w/w) of SAR247799, or control (CTRL) chow for 4 weeks. Age-matched lean ZSF1 (Le-ZSF1) rats were fed control chow. Echocardiography, telemetry, biochemical and histological analysis were performed to evaluate the effect of SAR247799.

**Results:** Echocardiography revealed that Ob-ZSF1 rats, in contrast to Le-ZSF1 rats, developed progressive diastolic dysfunction and cardiac hypertrophy with age. SAR247799 blunted the progression of diastolic dysfunction in young and old animals: in young animals E/e’ was evaluated at 21.8 ± 1.4 for Ob-ZSF1-CTRL, 19.5 ± 1.2 for Ob-ZSF1-SAR247799 p<0.01, and 19.5 ± 2.3 for Le-ZSF1-CTRL (median ± IQR). In old animals E/e’ was evaluated at 23.15 ± 4.45 for Ob-ZSF1-CTRL, 19.5 ± 5 for Ob-ZSF1-SAR247799 p<0.01, and 16.69 ± 1.7 for Le-ZSF1-CTRL, p<0.01 (median ± IQR). In old animals, SAR247799 reduced cardiac hypertrophy (mean ± SEM of heart weight/tibia length 0.053 ± 0.001 for Ob-ZSF1-CTRL vs 0.046 ± 0.002 for Ob-ZSF1-SAR247799 p<0.01, Le-ZSF1-CTRL 0.035 ± 0.001) and myocardial perivascular collagen content (p<0.001), independently of any changes in microvascular density. In young animals, SAR247799 improved endothelial function as assessed by the very low frequency bands of systolic blood pressure variability (mean ± SEM 67.8 ± 3.41 for Ob-ZSF1-CTRL 55.8 ± 4.27 or Ob-ZSF1-SAR247799, p<0.05 and 57.3 ± 1.82 Le-ZSF1-CTRL), independently of any modification of arterial blood pressure. In old animals, SAR247799 reduced urinary protein/creatinine ratio, an index of glomerular injury, (10.3 ± 0.621 vs 8.17 ± 0.231 for Ob-ZSF1-CTRL vs Ob-ZSF1-SAR247799, respectively, p<0.05 and 0.294 ± 0.029 for Le-ZSF1-CTRL, mean ± SEM) and the fractional excretion of electrolytes. Circulating lymphocytes were not decreased by SAR247799, confirming lack of S1P_1_ desensitization.

**Conclusions:** These experimental findings suggest that S1P_1_ activation with SAR247799 could improve LVH, cardiac diastolic and renal function in HFpEF patients.

## 1. Introduction

Heart failure with preserved ejection (HFpEF) affects approximately 50% of heart failure patients and, unlike heart failure with reduced ejection fraction, there are no drugs approved for HFpEF. Accompanied with the rise in life-expectancy, the prevalence of HFpEF continues to increase and its prognosis is bleak with a 5 year mortality rate over 50% (1, 2). So far clinical trials with approved drugs in heart failure with reduced ejection fraction have shown disappointing results on reducing mortality or hospitalization in HFpEF (3, 4). Thus, in order to bring forward effective treatments for HFpEF patients, there is a need to develop new drugs tailored to the pathophysiology of HFpEF *per se*. Left ventricular diastolic dysfunction plays a fundamental role in the pathophysiology of HFpEF. HFpEF is a complex syndrome in which patients present with multiple comorbidities including hypertension, diabetes, obesity, chronic kidney disease and atrial fibrillation (5). In addition to left ventricular diastolic dysfunction, these multiple risk factors result in left ventricular hypertrophy (LVH) that contributes to vascular dysfunction and increased myocardial stiffness.

Many studies have documented that endothelial dysfunction is prevalent in HFpEF patients, particularly in the microvasculature, and it is present in multiple vascular beds including the coronary, pulmonary, renal, or skeletal muscle circulations (6). Furthermore, established endothelial dysfunction predicts poor prognosis in HFpEF patients (6). According to the paradigm from Paulus and Tschope (7), the high prevalence of these comorbidities are responsible for a systemic proinflammatory state associated with endothelial dysfunction that causes reduced nitric oxide bioavailability and paracrine signaling changes in the cardiomyocyte, namely decreased cyclic guanosine monophosphate with subsequent titin hyper-phosphorylation and an increase in cardiomyocyte resting tension. Other consequences of endothelial dysfunction are reduced coronary flow reserve leading to myocardial ischemia, and transmigration of monocytes into the myocardium. All these changes are believed to cause concentric hypertrophy, cardiac stiffening, interstitial fibrosis and diastolic dysfunction (5).

Sphingosine 1-phosphate (S1P) is a lipid mediator secreted by endothelial cells, red blood cells and platelets, and exerts its effects through five receptors (S1P_1-5_) (8). Sphingosine-1-phosphate receptor 1 (S1P_1_) has the most prominent role on the endothelium, regulates vascular tone, inflammatory cell adhesion and migration, and endothelial cell barrier function (9). High S1P concentrations are found in the plasma of healthy individuals where it is carried primarily by HDL. HDL-bound S1P, acting through S1P_1_, limits vascular inflammation and atherosclerosis (10). Reduced plasma S1P and dysfunctional HDL are both found in a variety of patient populations with endothelial dysfunction. Plasma S1P is reduced in patients with systolic heart failure due to ischemic heart disease (11), in type 2 diabetic patients (12) as well as in rodent models of post-ischemic heart failure (13). Although several S1P_1_ modulators are either approved in multiple sclerosis (e.g. fingolimod, siponimod and ozanimod) or are in clinical development for immuno-inflammatory disorders, they were all developed as S1P_1_ functional antagonists that cause peripheral blood lymphopenia through S1P_1_ desensitization. In contrast, SAR247799 is a G-protein-biased S1P_1_ agonist, preferentially activating G-protein over β-arrestin and internalization signaling pathways and activates S1P_1_ without causing desensitization (14). SAR247799 thus exhibits endothelial-protective properties without reducing lymphocytes (15). SAR247799 improves the coronary microvascular hyperemic response in pigs, improves endothelial barrier integrity in a rat model of renal ischemia/reperfusion injury (15) and restores renal function and endothelial parameters in a rat model of type-2 diabetes (16). Moreover, SAR247799 demonstrates good safety and tolerability in healthy subjects (17) and improves endothelial function, as assessed by flow-mediated dilation in type-2 diabetic patients (16). The processes causing endothelial dysfunction are closely intertwined with cardiovascular disease and heart failure. Despite efforts to explore the effects of some endothelial-based therapies in the components of HFpEF, there is no confirmation at present that restoring endothelial function would provide clinical improvements in the progression of the components of HFpEF. We therefore set out to test whether 4-week oral treatment with the S1P_1_ agonist, SAR247799, a molecule that we have previously shown has endothelial protective effects in animals and humans (14–17) improves cardiac structural and functional parameters in a rat model of LVH and diastolic function. We used the diabetic Zucker fatty spontaneously hypertensive heart failure F1 hybrid (ZSF1) rat, carrying two separate leptin receptor mutations (fa and facp), which has been described as a genetically-hypertensive, obese and diabetic model of HFpEF (18, 19). ZSF1 rats have been described as a good model of HFpEF (18). Indeed, these rats present with the same risk factors found in HFpEF patients *i.e*. hypertension, obesity and type-2 diabetes and develop concentric LVH, diastolic dysfunction with preserved systolic function, pulmonary congestion and exercise intolerance, miming the features of human HFpEF (20). Although the ZSF1 rats do not reproduce the slow onset seen in patients with hypertension that lead to HFpEF development, this rat model may be argued to mimic the HFpEF patient in many scenarios (20), including the systemic proinflammatory state associated with endothelial dysfunction and was, thus, used in this study. As HFpEF patients are heterogeneous with respect to clinical criteria, we tested the effect of SAR247799 in two cohorts of obese ZSF1 rats that differed by age (termed young and old animals), and consequently the degree of diastolic dysfunction and hypertrophy.

## 2. Methods

### 2.1. Animals

Male obese ZSF1 (Ob-ZSF1) and Lean ZSF1 (Le-ZSF1) rats were purchased from Charles River (USA). Only male animals were considered in the present study, as at the time of the experiments only male ZSF1 rats were known to develop cardiac dysfunction. Two cohorts were obtained of different ages and considered for young and old animals protocols - as detailed below and in **Fig 1**. All animals were housed in groups of two in a climate-controlled environment (21-22°C, 55% humidity) with a 12:12 h light-dark cycle and provided with diet and water ad libitum. Control diet A04 (SAFE) was used from the arrival of the animals up to the starting of the experimental protocols. All animal studies were performed in accordance with the European Community standard on the care and use of laboratory animals and approved by the IACUC of Sanofi R&D. A lower number of lean animals was considered at the beginning of this study as lean rats were included as a comparison group to confirm the presence of LVH and diastolic dysfunction in ZSF1 rats and in order to follow the principle of reduction of 3Rs. At the end of each experiment, all animals were anesthetized under 3% isoflurane and euthanized by an intravenous injection of supersaturated KCl. All data were collected by blinded experimenters.

**Figure 1.**
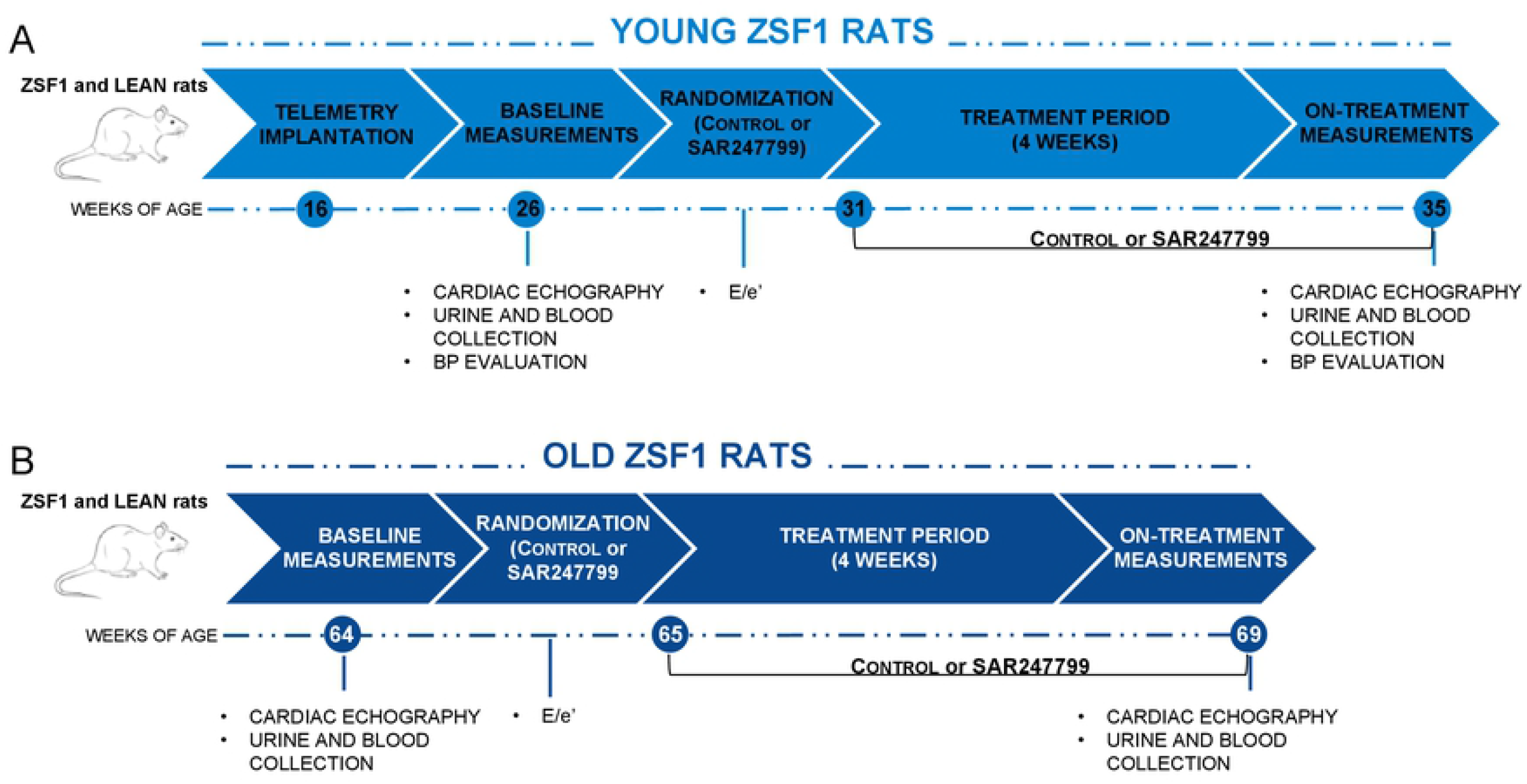
Overview of studies for young and old **ZSF1 rats**. Ob-ZSF1 rats were randomized on E/e’ parameter from doppler echocardiographic analysis. For young animal study, Le-ZSF1-CTRL N=9, Ob-ZSF1-CTRL N=11 and Ob-ZSF1-SAR247799 N=11. For old animals study Le-ZSF1-CTRL N=9, Ob-ZSF1-CTRL N=7 and Ob-ZSF1-SAR247799 N=7; N= number of rats.

### 2.2. In vivo experimental protocols

#### 2.2.1. Young ZSF1 rats

10-12-week-old rats (22 Ob-ZSF1 and 9 Le-ZSF1 rats) were purchased and, 4-weeks after arrival, were implanted with a telemetry device (Data Sciences International^™^), as described below. At 26 weeks of age, rats underwent a baseline echocardiography, blood and urine collection. Ob-ZSF1 rats were randomized based on the diastolic function parameter E/e’ to control chow (Ob-ZSF1-CTRL N=11) or to chow formulated with 0.025 % (w/w) SAR247799 (Ob-ZSF1-SAR247799 N=11), (SSNIFF Spezialdiäten GmbH). Le-ZSF1 rats were maintained on control chow. Post treatment assessments were made 4 weeks later by echocardiography, as well as blood and urine collection.

#### 2.2.2. Old ZSF1 rats

5-7-week-old rats (22 Ob-ZSF1 and 9 Le-ZSF1, rats) were purchased and kept in the animal facilities up to 64 weeks of age, when baseline echocardiography was performed, and samples of blood and urine collected on the animals alive (14 Ob-ZSF1 and 9 Le-ZSF1). Ob-ZSF1 rats were randomized based on the E/e’ value to a control chow diet (Ob-ZSF1-CTRL N=7) or to a diet with chow supplemented with 0.025 % (w/w) SAR247799 (Ob-ZSF1-SAR247799 N=7), (SSNIFF Spezialdiäten GmbH). Le-ZSF1 rats were maintained on control chow. Post-treatment echocardiography was performed 4 weeks after randomization, at which point urine and blood were also collected. Histological analyses were performed on cardiac tissues as detailed below.

### 2.3. Telemetry measurements

In order to evaluate blood pressure and systolic blood pressure (SBP) variability (frequency-domain analysis), rats included in the young animal group were equipped with a telemetry device. Rats were anesthetized with 4.5% isoflurane, which was then progressively lowered and maintained at 2% until the end of the surgery. Rats were placed on thermostatically-controlled heating pads, and body temperature was maintained at 37°C. Animals were then administered several drugs: a central analgesic (fentanyl at 10 μg/kg, SC), a mixture of local analgesics (lidocaine at 2 mg/kg and bupivacaine at 2 mg/kg, on surgery site, SC), an anti-inflammatory (carprofen at 4 mg/kg, SC) and an antibiotic (long acting terramycine at 30 mg/kg IM the day of surgery and 3 days later). Rats were implanted with HD-S11 devices, placed in the abdomen, for BP measurement, which was recorded through a catheter inserted into the abdominal aorta. At the end of the surgery, after recovery from anesthesia in a temperature-controlled ventilated room, rats received buprenorphine (10 μg/kg, SC) and were housed in separate cages for 10 days, during which they received carprofen and buprenorphine (4 mg/kg and 10 μg/kg respectively, SC) daily on the first 2 days. Rats were subsequently housed in groups of two per cage.

Telemetry device data were collected using Hem 4.3 acquisition software (Notocord Systems) connected to DSI’s telemetry device. At baseline and 4 weeks after treatment initiation, diastolic and systolic blood pressures were recorded continuously for 24 hours starting at 11 am. For the frequency domain analysis, SBP data points were re-sampled at 8 Hz. Periods of 512 points without erratic fluctuations (due to interruptions or interferences derived from animal movements) were visually identified and used for the calculation of the SBP variability.

Fast Fourier transformation was performed, and power spectra of each period were extracted using Hem 4.3. The software was set to determine very low frequencies (VLF: 0.02-0.2 Hz), low frequencies (LF 0.2-0.6 Hz) and high frequencies (HF: 0.6 to 4 Hz) spectral powers. Spectral powers are the area under the curve of the power spectra in the respective frequency bands. VLF are expressed in normalized units (n.u.) (i.e. the percentage of the VLF power with respect to the sum of the VLF, LH and HF powers) (21).

### 2.4. Echocardiography

To evaluate diastolic and systolic function and left ventricular geometry, transthoracic cardiac echography was performed. Rats were anesthetized in an induction cage with 3% isoflurane and then maintained to steady-state sedation level with 2% isoflurane by the nose connected to the anesthesia system. The desired depth of anesthesia was confirmed in animals by the absence of reflex after toe pinch. Rats were placed in a decubitus position on a warming pad, the chest was shaved, and hair removal cream was used to remove the remaining body hairs. A gel (Ocry-gel, TVM) was applied to both eyes to prevent drying of the sclera and a lubricated rectal probe was inserted to continuously monitor body temperature in order to maintain it in a range of 37°C ± 0.5. ECG was recorded from suitable electrodes connected to Affiniti 50 (Philips). A layer of preheated gel was applied over the chest. Two-dimensional, M-mode and pulse wave Doppler were obtained from a 15-MHz ultrasound probe, while tissue Doppler at mitral annulus level was recorded from a 12-MHz ultrasound probe. Video were recorded and then analyzed off-line in Q-Station (Philips). All measures were taken from five cardiac cycles and then averaged.

### 2.5. Plasma and urinary evaluations

18-hour urine samples were collected in metabolic cages (Tecniplast). Urinary protein, creatinine, chloride and sodium concentration were measured using a PENTRA 400 analyzer (Horiba). Blood samples were collected in K_2_-EDTA tubes from the caudal vein in animals under 1.5-2% isoflurane anesthesia. Blood lymphocyte count was measured by MS9-5 (Melet Schloesing) in freshly collected blood, and then blood samples were immediately centrifuged at 4°C/10 min/10,000g before storing plasma at −80°C. SAR247799 concentration was measured in 4-week post-treatment plasma samples by a qualified liquid chromatography tandem mass spectrometry method using a Shimadzu chromatographic system coupled to an API 4000 mass spectrometer (Sciex). Plasma fructosamine, creatinine, chloride and sodium concentration were measured using a PENTRA 400 analyzer. Fractional excretion of sodium (FENa) and chloride (FECl) were then calculated by (urinary sodium / plasma sodium)*(plasma creatinine/ urinary creatinine)*100 and (urinary chloride / plasma chloride)*(plasma creatinine/ urinary creatinine)*100. Daily food intake was recorded weekly in young and old animals.

### 2.6. Collagen content and microvascular quantification

Transversal 5-μm-thick sections were cut from formalin-fixed and paraffin-embedded heart slices and mounted onto microscope slides. Masson’s trichrome stain was used for collagen content quantification. Immunohistochemical staining of vessels was carried out using the Ventana Discovery XT automated system with recommended reagents (Ventana Medical Systems Inc). Tissue slides were dewaxed and pretreated with cell-conditioning buffer CC1 at 95°C for 48 minutes. An anti-CD31 antibody (ab182981, Abcam) was used at 5 μg/mL final concentration for 1 hour at 24°C following by goat anti-rabbit IgG biotin conjugate used as secondary antibody at 1-200 final dilution (BA-1000, Vector Laboratories). DAB Map including 3,3-Diaminobenzidine chromogen was applied to the slides for histological staining.

Digital images of Masson’s trichrome stained sections were captured at 20X magnification using Aperio AT2 scanner (Aperio Leica) and analyzed with the HALO image analysis platform (Indica labs). A tissue classifier add-on was applied to identify large vessels and myocardial tissue using machine learning with random forest algorithm based on color texture and contextual features. A positive pixel count was applied to quantify the collagen ratio with HALO area quantification algorithm in the interstitial myocardial tissue and large vessels. Microvasculature was quantified on CD31-stained slides. Five regions of interest (2 mm^2^ each) were selected in transversal sectioned myofibers of the left ventricle. Microvessel density (with a vessel area ≤5000 mm^2^) was measured using ImageScope microvessels algorithm (Aperio Leica) and expressed as microvascular density/mm^2^.

### 2.7. Data and statistical analysis

The data and statistical analysis comply with the recommendations on experimental design and analysis in pharmacology (22). In each study (young and old animals groups), statistical analyses were performed at baseline on each parameter to compare Le-ZSF1 rats to the 2 groups of Ob-ZSF1 rats prior to randomization. Then, after 4-weeks of treatment, the pathology in the Ob-ZSF1-CTRL group was evaluated by statistical comparison to the Le-ZSF1-CTRL group. The effect of SAR247799 treatment was then compared to control chow treatment in Ob-ZSF1 rats. For each analysis, when the hypothesis of normality was fulfilled, a Student test was performed, otherwise a Wilcoxon test was performed. p<0.05 was considered significant. The approach to perform distinct analyses before and after treatment was chosen in order to perform the statistical analyses taken into account exactly the same experimental conditions, and avoid impact of these on the results. Moreover, a distinct statistical analysis at baseline allows to take into account all the ZSF1 animals to evaluate the pathology effect with more power. Post treatment two models were performed to answer two distinct objectives: evaluation of the pathology and then of the treatment effect. One model by objective was chosen to gain power in case of heteroscedasticity. Data are expressed as mean ± SEM or median ± IQR (interquartile range). All statistical analyses were performed with SAS^®^ version 9.4 for Windows 7. The authors confirm that the data supporting the findings of this study are available within the article.

## 3. Results

### 3.1. Characteristics of young and old obese ZSF1 rats

Compared to age-matched Le-ZSF1 rats, 26-week-old Ob-ZSF1 rats developed slight cardiac hypertrophy (significantly increased left ventricular posterior wall at end-diastole, LVPWd, thickness) and diastolic dysfunction (significantly increased E/e’ ratio), together with a preserved ejection fraction % **(Supplementary Table I)**. 64-week-old Ob-ZSF1 rats, compared to their age-matched Le-ZSF1, displayed diastolic dysfunction (significantly increased E/e’) and significantly more pronounced left ventricular hypertrophy, with both thickened inter-ventricular septum at end-diastole (IVSd) and LVPWd. In both studies, Ob-ZSF1 rats displayed a significantly increased stroke volume and similar cardiac index compared to age-matched Le-ZSF1 rats. Relative to age-matched Le-ZSF1 rats, chambers volumes were preserved in the young animals, while end-diastolic volume increased significantly in the old animals.

### 3.2. SAR247799 delays progression of diastolic dysfunction in young and old obese ZSF1 rats

The effect of SAR47799 on diastolic function was assessed by measuring the E/e’ ratio, an index of ventricular filling pressure, by echocardiography. SAR247799 consistently limited the deterioration of diastolic function, as shown by a significant reduction of the E/e’ ratio **(Fig 2 A)** in both young and old ZSF1 rats. This reduced E/e’ was mainly due to restored mitral annular velocity during early diastole (e’) and to lesser extent to improved mitral flow velocity during early diastole (E wave) **(Fig 2 B & C)**. E/A ratio measured in young animals did not reach statistical significance difference in the model nor with the compound (**Supplementary Fig 2**). SAR247799 did not modify indices of systolic function including ejection fraction % **(Fig 2 D)**, the end-diastolic volume and the end-systolic volume **(Fig 2 E & F)**, the stroke volume or the cardiac index **(Fig 2 G & H)** in either young or old ZSF1 rats.

**Figure 2.**
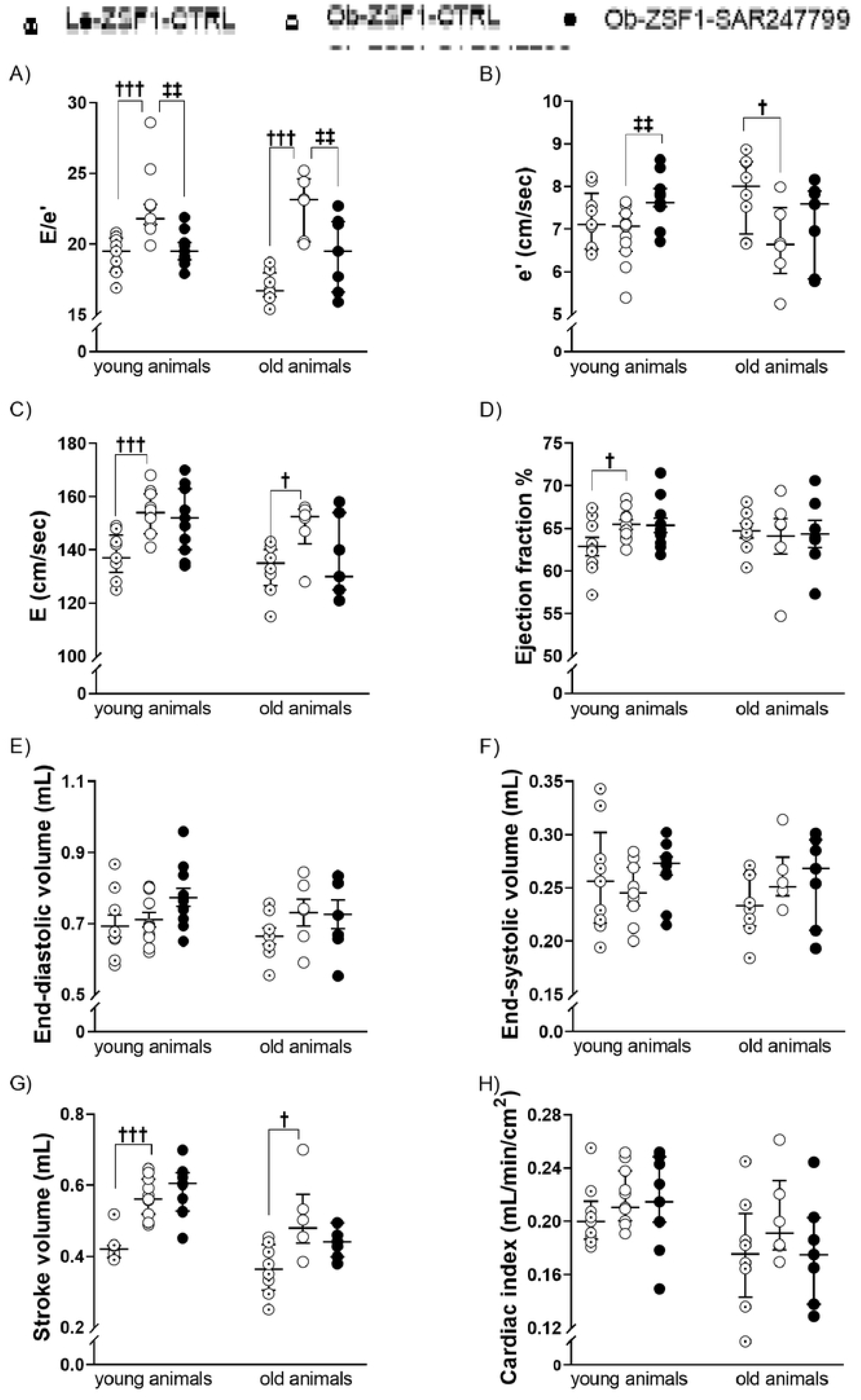
Cardiac diastolic and systolic function in young and old ZSF1 rats after 4 weeks of treatment. Diastolic function was evaluated by (A) E/e’ ratio (B) septal peak e’ wave velocity at mitral annulus level and (C) mitral valve peak E wave velocity. Cardiac systolic function was assessed by (D) ejection fraction, (E) end-diastolic volume, (F) end-systolic volume, (G) stroke volume and (H) cardiac index. Data are expressed mean ± SEM, except E, e’, E/e’, end-systolic volume, stroke volume and which are expressed as median ± IQR. **†**p<0.05, **†††** p<0.001: comparison of Le-ZSF1-CTRL to Ob-ZSF1-CTRL using a Student t-test except for E/e’ and stroke volume for young animals and for E wave for old animals for which a Wilcoxon test was performed. **‡**p<0.05, **‡‡**p<0.01: p-values obtained from the comparison between Ob-ZSF1-CTRL and Ob-ZSF1-SAR247799 using a Student t-test except for E/e’ for young animals and for E wave for old animals for which a Wilcoxon test was performed. For young animals, Le-ZSF1-CTRL N=9, Ob-ZSF1-CTRL N=11 and Ob-ZSF1-SAR247799 N=11. For old animals Le-ZSF1-CTRL N=8-9, Ob-ZSF1-CTRL N=6-7 and Ob-ZSF1-SAR247799 N=7; N= number of rats. A lower number of n values is due to technical issue in echocardiography data acquisition.

### 3.3. SAR247799 reduced cardiac hypertrophy in old ZSF1 rats

The degree of cardiac hypertrophy was quantified in the two models by heart and heart chamber weights as well as by chamber morphometric parameters assessed by echocardiography **(Fig 3)**. Compared to age-matched Le-ZSF1-CTRL rats, Ob-ZSF1-CTRL rats displayed increase in total cardiac hypertrophy (evaluated as heart weight/tibia length), left atrial weight/tibia length, right atrial weight/tibia length, IVSd, inter-ventricular septum at end-systole (IVSs), LVPWd and left ventricular posterior wall at end-systole (LVPWs) **(Fig 3 A-G),** more marked in old than in young animals. In old animals, SAR247799 significantly reduced heart weight/tibia length **(Fig 3 A)**, right atrial weight/tibia length **(Fig 3 C)**, inter-ventricular septum at end-systole (IVSs) **(Fig 3 E)** and LVPWs **(Fig 3 G)** and showed a trend to reduce left atrial weight/tibia length **(Fig 3 B)**, IVSd **(Fig 3 D)** and LVPWd **(Fig 3 F)**. In young animals, SAR247799 significantly reduced right atrial weight/tibia length **(Fig 3 C)** and IVSs **(Fig 3 E)**, while have little effect on the other parameters. Overall SAR247799 reduced cardiac hypertrophy in both young and old animals groups, but the anti-hypertrophic effect was more pronounced in old animals which exhibited more advanced hypertrophy. Representative echocardiographic images for E wave, e’ wave and left ventricular thickness from TM-mode are shown in Supplementary Fig 2 (A-C) for young animals and (D-F) for old animals.

**Figure 3.**
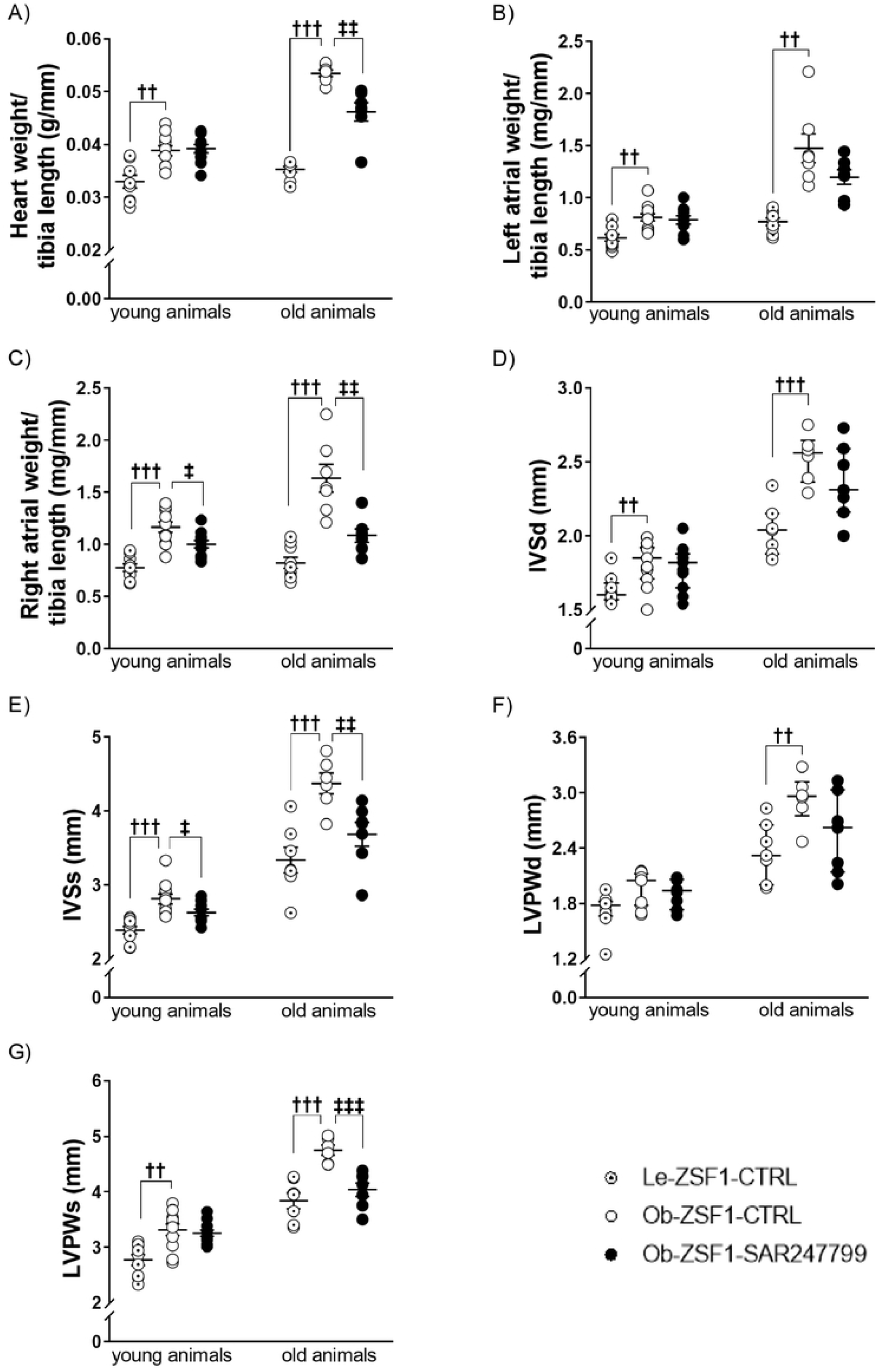
Cardiac hypertrophy in young and old ZSF1 rats after 4 weeks of treatment. **(A)** Heart weight/tibia length, **(B)** left atrial weight/tibia length, **(C)** right atrial weight/tibia length, **(D)** interventricular septum thickness at end-diastole (IVSd), **(E)** interventricular septum thickness at end-systole (IVSs), **(F)** left ventricular posterior wall thickness at end-diastole (LVPWd), **(G)** left ventricular posterior wall thickness at end-systole (LVPWs). Data expressed as mean ± SEM, except IVSd and LVPWd which are expressed as median ± IQR. **††** p<0.01, **†††** p<0.001: comparison of Le-ZSF1-CTRL to Ob-ZSF1-CTRL using a Student t-test, except LVPWd (young animals) for which a Wilcoxon test was performed. **‡**p<0.05, **‡‡**p<0.01, **‡‡‡**p<0.001: comparison between Ob-ZSF1-CTRL and Ob-ZSF1-SAR247799 using a Student t-test, except LVPWd (young animals) for which a Wilcoxon test was performed. For young animals, Le-ZSF1-CTRL N=9, Ob-ZSF1-CTRL N=11 and Ob-ZSF1-SAR247799 N=10-11. For old animals Le-ZSF1-CTRL N=8-9, Ob-ZSF1-CTRL N=6-7 and Ob-ZSF1-SAR247799 N=7; N= number of rats. A lower number of n values is due to technical issue in echocardiography data acquisition.

### 3.4. SAR247799 reduced cardiac fibrosis in old ZSF1 rats independent of microvascular density

Compared to Le-ZSF1-CTRL rats, Ob-ZSF1-CTRL rats displayed an increase in perivascular collagen in the old ZSF1 rats and similar levels of interstitial collagen **(Fig 4 A-B-C)**. SAR247799 significantly reduced perivascular collagen and tended to decrease interstitial collagen. Microvascular density in the left ventricle was decreased in Ob-ZSF1-CTRL rats compared to Le-ZSF1-CTRL. SAR247799 did not restore cardiac microvascular density over the 4-week treatment period **(Fig 4 D)**.

**Figure 4.**
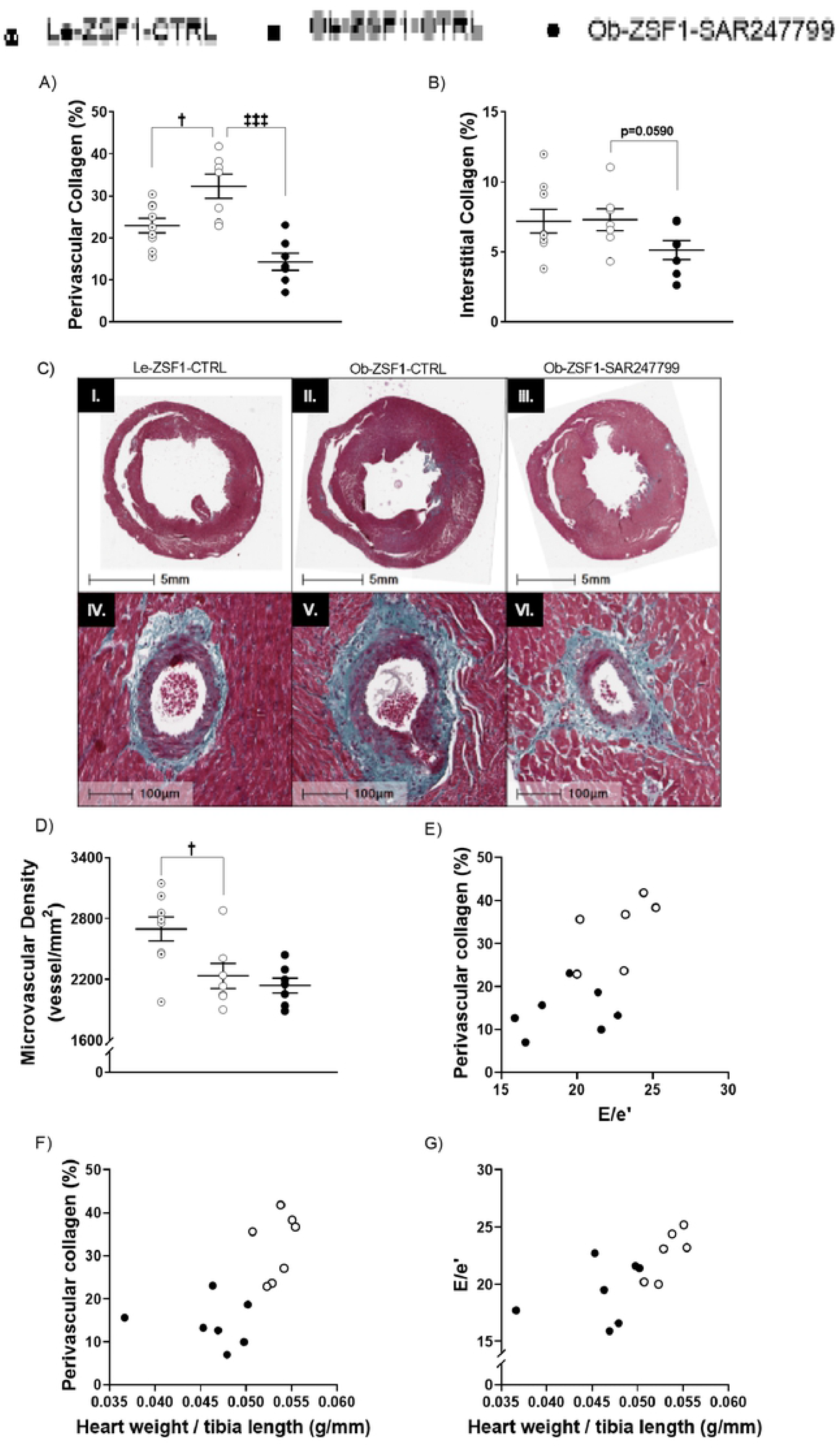
Cardiac fibrosis and microvascular density after 4 weeks of treatment and relationships between diastolic dysfunction, cardiac hypertrophy and fibrosis in old ZSF1 rats. **(A)** perivascular collagen, **(B)** interstitial collagen, **(C)** representative histological images of cardiac fibrosis in Le-ZSF1-CTRL (I.), Ob-ZSF1-CTRL (II.) andOb-ZSF1-SAR247799 (III.), scale bar 5 mm and corresponding perivascular fibrosis (IV., V. and V.,) stained in green by Masson trichrome staining, scale bar, 100 μm. **(D)** microvascular density. Scatter plot showing relationships between **(E)** perivascular collagen and diastolic dysfunction (E/e’), **(F)** perivascular collagen and heart weight/ tibia length (normalized cardiac hypertrophy), **(G)** diastolic dysfunction and heart weight/ tibia length. For **(A, B & D)** data presented as mean ± SEM. **†**p<0.05: p-values obtained to compare Le-ZSF1-CTRL to Ob-ZSF1-CTRL using a Student t-test. **‡‡‡**p<0.001: p-values obtained from the comparison between Ob-ZSF1-CTRL and Ob-ZSF1-SAR247799 using a Student t-test. Le-ZSF1-CTRL N=9, Ob-ZSF1-CTRL N=7 and Ob-ZSF1-SAR247799 N=7; N= number of rats.

### 3.5. SAR247799 concomitantly ameliorated diastolic function, fibrosis and hypertrophy indices in old ZSF1 rats

Relationships between the effects of SAR247799 on diastolic function, hypertrophy and fibrosis, were illustrated with scatter plots for the respective parameters, E/e’, heart weight/tibia length and perivascular collagen. SAR247799-treated rats displayed clear separated values when compared to Ob-ZSF1-CTRL rats for the relationships involving perivascular collagen with diastolic dysfunction (E/e’) **(Fig 4 E)**, perivascular collagen with normalized cardiac hypertrophy (heart weight/tibia length) **(Fig 4 F)** and diastolic dysfunction with normalized cardiac hypertrophy **(Fig 4 G)**. Overall, SAR247799 improved diastolic function, hypertrophy and fibrosis concomitantly.

### 3.6. SAR247799 improved endothelial function without modifying blood pressure

Vascular effects of SAR247799 were investigated in young ZSF1 rats, previously implanted with a transducer allowing pressure measurement of the abdominal aorta. At baseline, Ob-ZSF1 rats displayed similar systolic and an increased diastolic blood pressure compared to Le-ZSF1 rats. SAR247799 did not modify systolic or diastolic blood pressure (**Fig 5 A & B**).

**Figure 5.**
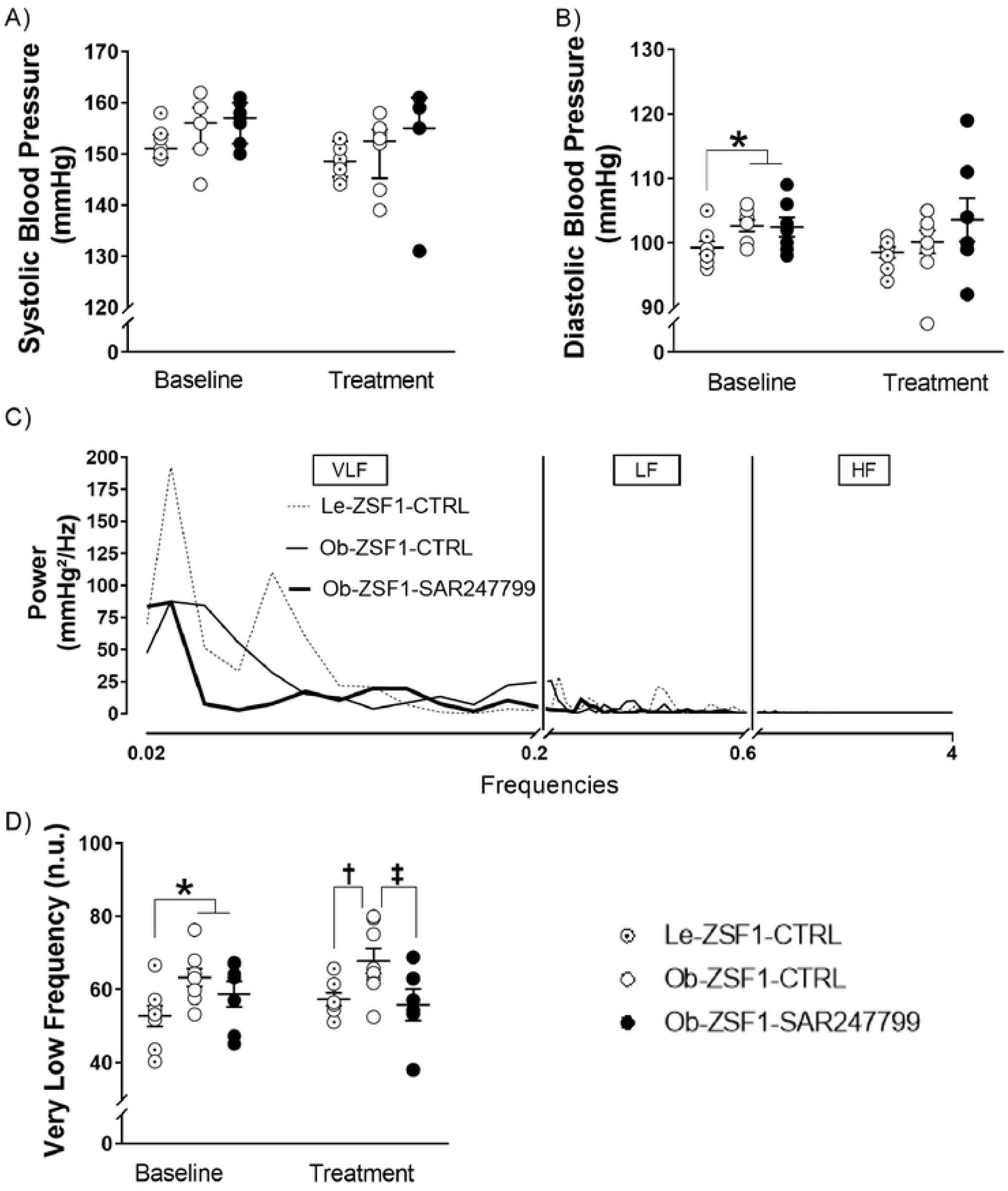
Blood pressure and blood pressure variability after 4 weeks of treatment in young animals. **A)** systolic blood pressure (SBP), **(B)** diastolic blood pressure, **(C)** representative spectra of systolic blood pressure variability showing very low frequency (VLF), low frequency (LF) and high frequency (HF) bands. **(D)** very low frequency (n.u.) bands from spectral analysis of SBP variability. Data presented as mean ± SEM, except SBP who is presented as median ± IQR. **\***p<0.05, **\*\***p<0.01: p-values obtained to compare Le-ZSF1-CTRL to Ob-ZSF1 (Ob-ZSF1-CTRL and Ob-ZSF1-SAR247799 pooled) using a Student t-test. A Student t-test was also performed to compare Ob-ZSF1-CTRL to Ob-ZSF1-SAR247799 at baseline. No significant difference was observed between the two randomized groups at baseline for any of the parameters. **†**p<0.05: p-values obtained to compare Le-ZSF1-CTRL to Ob-ZSF1-CTRL using a Student t-test. **‡**p<0.05: p-values obtained from the comparison between Ob-ZSF1-CTRL and Ob-ZSF1-SAR247799 using a Student t-test except for SBP for which a Wilcoxon test was performed. For young animals, Le-ZSF1-CTRL N=8, Ob-ZSF1-CTRL N=8 and Ob-ZSF1-SAR247799 N=7; N= number of rats. The lower number of n values is due to technical issue in telemetry data acquisition.

We further investigated systolic blood pressure variability by spectral methods and computed VLF, LF and HF (**Fig 5 C**) bands of the power of the spectrum. Frequency analysis of blood pressure tracing using Fourier transformation consist of a quantitative measurement of blood pressure variability at distinctive frequency bandwidths that have been associated with blood pressure control mechanisms. Indeed, vasoactive agents may affect variability without changing absolute blood pressure (21, 23). In rats, HF bands of SBP variability have been associated with β-adrenoceptor-mediated cardiac sympathetic function or to the mechanical effect of respiration (24), LF bands to vascular sympathetic activity, and VLF bands to endothelial-derived nitric oxide action (21). Thereby, the integral of VLF (0.02-0.2 Hz) is inversely associated with the activity of the endothelial nitric oxide system (21). In our study VLF was significantly higher in Ob-ZSF1-CTRL rats compared to Le-ZSF1-CTRL rats and SAR247799 significantly reduced this parameter, implying an improvement in endothelial function (Fig 5 D).

### 3.7. SAR247799 improved renal function in old ZSF1 rats

Renal function parameters in Ob-ZSF1 and Le-ZSF1 rats in young and old ZSF1 rats, at baseline and after 4 weeks of treatment are summarized in **Table I**. In old animals, Ob-ZSF1 rats displayed a decrease in urinary function compared to Le-ZSF1 littermates, as demonstrated by significant increases in protein/creatinine ratio, FENa and FECl. Urinary flow was elevated at baseline in Ob-ZSF1-CTRL rats compared to Le-ZSF1, and to a lesser extent 4 weeks later, consistent with the beginning of end-stage renal failure. Four-week treatment with SAR247799 significantly decreased urinary protein/creatinine ratio and FECl, indicating that renal filtration was ameliorated. In young animals, SAR247799 did not modify renal parameters in Ob-ZSF1 rats that started to display renal dysfunction (increased urinary flow, urinary protein/creatinine ratio and FENa).

**TABLE I:**
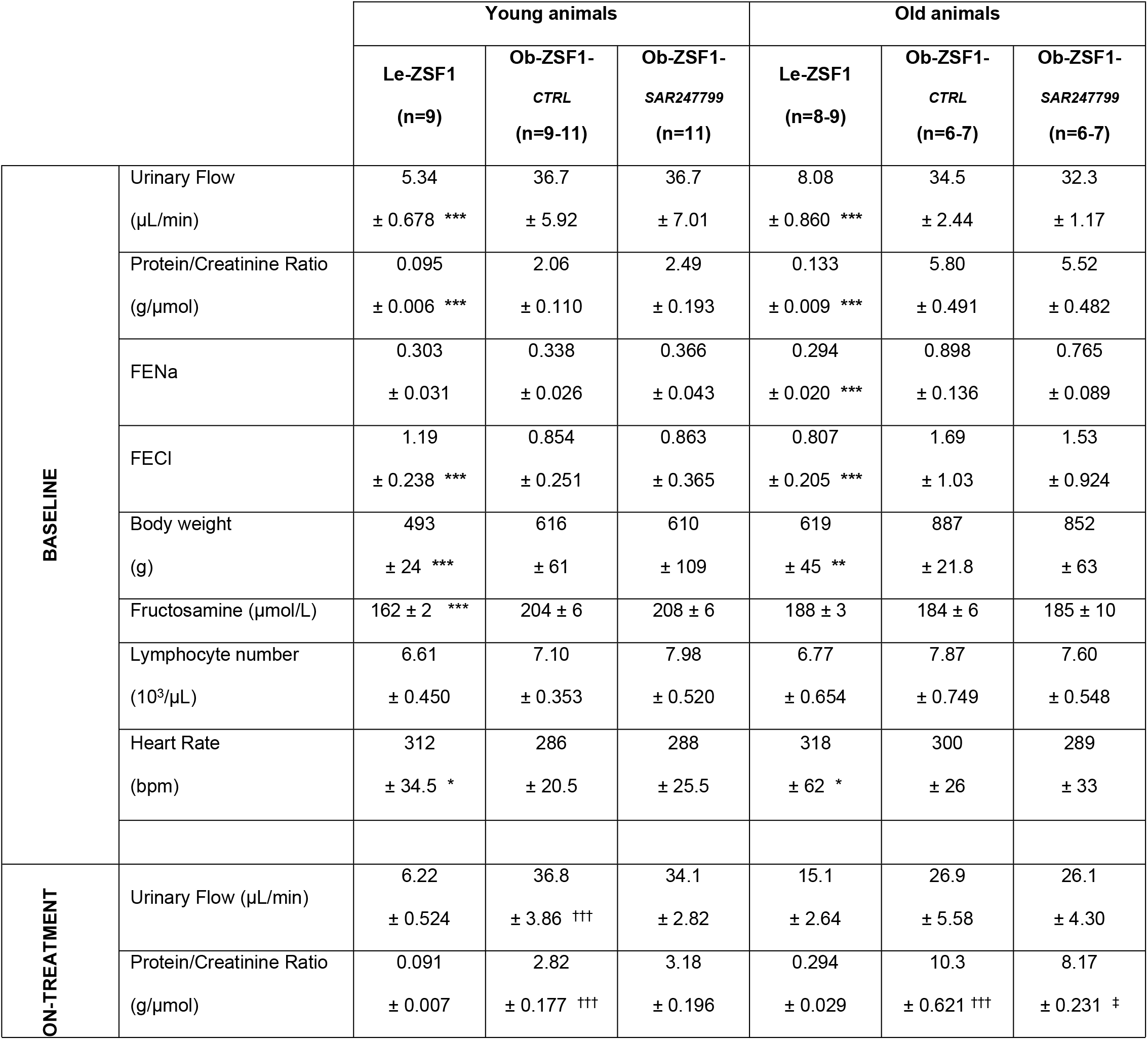
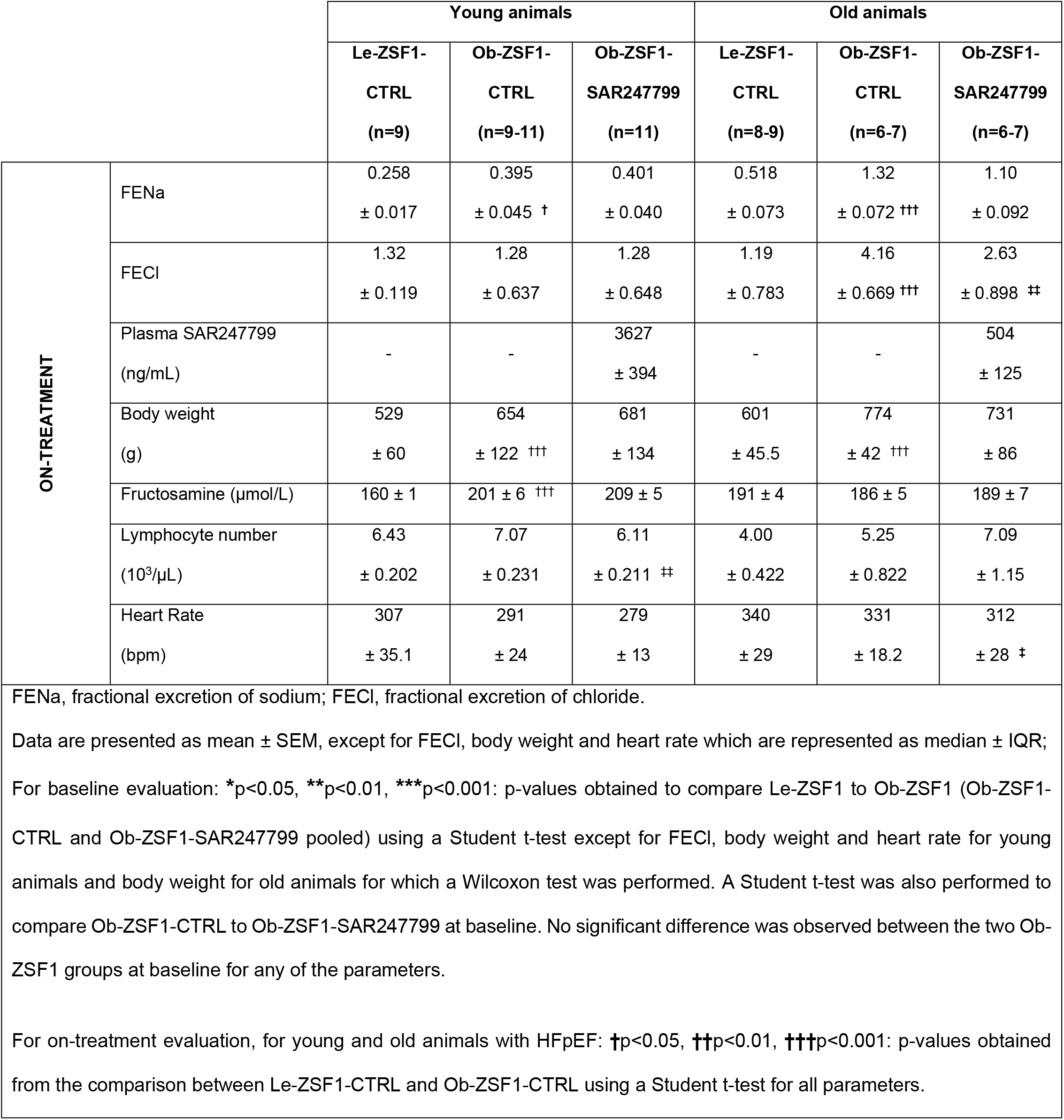
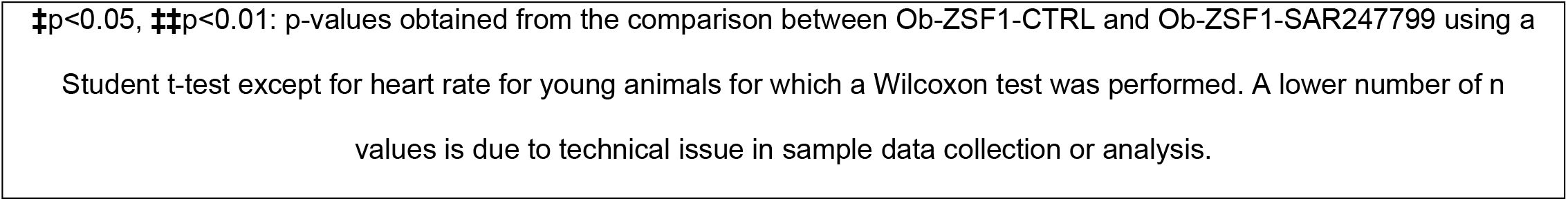
Renal function and physiological parameters at baseline and after 4 weeks of treatment

### 3.8. Effects of SAR247799 on physiological parameters

SAR247799 was incorporated in the chow at the dose of 250 mg/kg as a previous study from our laboratory demonstrated the effect of this dose in endothelial protection in another rat model of metabolic dysfunction (16). In this study, the plasma concentration of SAR247799 after 4 weeks of treatment was 7-fold higher in young compared to the old ZSF1 rats. However, daily food intake was greater in young than in old animals (**Supplementary Table II**).

The effect of SAR247799 on lymphocytes (a marker of S1P_1_ desensitization) and heart rate (a marker of cardiac S1P_1_ activation) were determined following 4-week treatment, alongside measures of body weight and compound exposure (**Table I**). Following the 4-week treatment periods in young and old animals, blood lymphocyte counts were similar between Ob-ZSF1-CTRL and Ob-ZSF1-SAR247799 groups, indicating that the dosing regimens did not cause S1P_1_ desensitization. Moreover, SAR247799 reduced heart rate by 4% (NS) and 6% (significant), in young and old ZSF1 rats, respectively, compared to the Ob-ZSF1-CTRL group. Ob-ZSF1 rats (both models) were obese compared to age-matched Le-ZSF1 rats, and SAR247799 treatment did not modify body weight. Plasma fructosamine was increased in Ob-ZSF1 rats compared to Le-ZSF1 in young but not in old animals and these levels were not affected by SAR247799 treatment (Table I).

## 4. Discussion

The present study provides the first evidence that SAR247799 confers cardio-protective effects in experimental models of LVH and diastolic dysfunction, due to S1P_1_ activation. We demonstrated that 4-week treatment with the G-protein-biased S1P_1_ agonist, SAR247799, blunted the progression of diastolic dysfunction, and reduced cardiac hypertrophy, right atrial enlargement and perivascular cardiac fibrosis. SAR247799 also had renal-protective effects and did not modify arterial blood pressure. The protective effects of SAR247799 were under conditions where there was no S1P_1_ desensitization (no lymphocyte reduction) and there was an improvement in endothelial function, in line with the proposed mechanism of action.

The Ob-ZSF1 rat mimicked closely the features found in HFpEF patients, i.e. LVH and diastolic dysfunction. HFpEF patients are characterized by having a preserved ejection fraction and impaired diastolic function, as was seen in the Ob-ZSF1 rats (18, 19). Hypertension, obesity and diabetes are significant human risk factors affecting the progression to HFpEF and were characteristic features of Ob-ZSF1 rats. Diabetes and obesity contributed to the HFpEF phenotype as Ob-ZSF1 rats had more marked pathological changes compared to their respective non-diabetic age-matched Le-ZSF1 rats which displayed hypertension alone. Old age is an important risk factor for the LVH and diastolic dysfunction progression. The inclusion of the older Ob-ZSF1 cohort (65 weeks of age) revealed higher degrees of concentric cardiac hypertrophy and atrial remodeling compared to the younger Ob-ZSF1 cohort (31 weeks of age), features described in more than 50% of HFpEF patients (5). Young ZSF1 displayed diastolic dysfunction with minimal left ventricular hypertrophy while in old ZSF1 left ventricular hypertrophy was highly present with diastolic dysfunction. This evidence suggested that left ventricular hypertrophy is establishing with ageing and consequently to diastolic dysfunction and uncontrolled blood pressure. Myocardial fibrosis usually presents as perivascular and fine interstitial reactive fibrosis and is more evident in advanced HFpEF (19, 25), thus was here evaluated in old ZSF1 rats. Myocardial fibrosis is known to play a major role in the development of LVH, diastolic dysfunction and thus HFpEF. In patients, myocardial fibrosis is associated with death and hospitalization for heart failure in proportion to the severity of fibrosis (25). In our old Ob-ZSF1, a good amount of perivascular but not interstitial fibrosis was found. Fibrotic tissue deposition in perivascular spaces impairs oxygen supply to the cardiomyocyte. Indeed, perivascular fibrosis is inversely correlated with coronary flow reserve in heart failure patients (26). Importantly, our results showed that SAR247799 had cardio-protective effects in both young and old ZSF1 rats, suggesting its potential utility in a broad range of HFpEF patients.

The protective effects of SAR247799 in Ob-ZSF1 rats could be attributed to its S1P_1_-mediated endothelial-protective properties. Previous studies demonstrate that S1P_1_ activation improves outcomes after myocardial or cerebral ischemia/reperfusion injury (27) and that cardiac overexpression of S1P_1_ improves left ventricular contractility and relaxation in rats undergoing myocardial infarction (13), while endothelial S1P_1_ deficient mice display worse outcomes (28). Indeed, we showed in the present study in Ob-ZSF1 rats, that SAR247799, through its mechanism of action on endothelial function, reduced VLF bands of systolic blood pressure variability, an index of endothelial nitric oxide pathway activation (23). Similarly, our previous work shows that SAR247799 improves coronary microvascular reserve in a pig model of ischemic injury (15) and endothelial and renal function in diabetic rats (16). Furthermore, SAR247799 improves flow-mediated dilation in type-2 diabetes patients (16). Collectively, SAR247799 has endothelial-protective properties consistent with a role for the cardio-protective effects seen in experimental LVH and diastolic dysfunction. As active relaxation is an energy-dependent phase of diastole, vulnerable to ischemia, SAR247799’s ability to improve coronary microvascular perfusion (15) could explain the improvements in diastolic function. Furthermore, longstanding myocardial ischemia and inflammation induce myocardial hypertrophy and fibrosis, resulting in decreased left ventricular compliance. SAR247799’s ability to improve myocardial perfusion and reduce vascular inflammation (15) could therefore prevent further deterioration of the extracellular matrix and fibrosis. Endothelial cells also have an important role in paracrine signaling to the cardiomyocyte through cyclic guanosine monophosphate activation and regulating titin hyperphosphorylation and subsequent cardiac hypertrophy (19). Given the improvements in hypertrophy seen in the model, the anti-hypertrophic effects of SAR247799 could have been mediated by restoring this paracrine signaling. Moreover, in a previous study in Zucker Diabetic Fatty (ZDF) rats, SAR247799 was able to reduce not only plasma levels of adhesion molecules such as sICAM-1 and sE-selectin but also CRP, an inflammatory marker (16). For these reasons, we hypothesize that SAR247799 reduced cardiac hypertrophy and fibrosis by reducing inflammation and/or improving coronary and peripheral endothelial cell function. The anti-hypertropic and anti-fibrotic effects in the heart shown herein with SAR247799 are consistent with a recent study in which endothelial-specific S1P_1_ knockdown promoted cardiac hypertrophy and fibrosis in a pressure overload model (29). Furthermore, the protective effects seen with SAR247799 in the old ZSF1 rats (improved diastolic function, reduced cardiac hypertrophy, reduced cardiac fibrosis and improved endothelial and renal function) were at plasma concentrations (mean+SD 504+125 ng/ml) that were in a similar range to concentrations causing endothelial-protective effects in humans, rats and pigs, as well as on endothelial cells (see **Table II**). There is evidence for microvascular rarefaction in HFpEF patients, as coronary microvascular density is reduced in cardiac autopsies (7, 30). Similarly, our experimental **ZSF1 rat model** had reduced microvascular density in old animals. Although S1P_1_ activation plays an essential role in angiogenesis and vasculogenesis, 4-week treatment with SAR247799 did not improve microvascular density. Consequently, the significant improvements in cardiac parameters found with SAR247799, over this timeframe, cannot be explained by changes in microvessel density, but may be more attributed to structural and functional modifications of the vessels. As 12-week S1P_1_ gene therapy in ischemic rats has been shown to improve coronary microvessel density (13), the potential for higher efficacy with longer treatment durations of SAR247799 warrants consideration.

**TABLE II.** Comparison of the protective effects of SAR247799 in Old obese ZSF1 rats with effects seen in prior experimental systems

| Experimental system | Protective effects shown | Plasma concentration associated with effects (ng/ml)<br>Mean±SD | Reference |
| --- | --- | --- | --- |
| Old Ob-ZSF1 rats | Improved diastolic function<br>Reduced cardiac hypertrophy<br>Reduced cardiac fibrosis<br>Improved endothelial function (VLF)<br>Reduced urinary protein/creatinine ratio<br>Reduced fractional excretion of electrolytes | 504±125 | Current study |
| ZDF rat (intermediate dose strength) | Reduced plasma endothelial biomarkers (sICAM1, sE-Selectin, VWF, nitrite/nitrate)<br>Reduced urinary protein/creatinine ratio<br>Reduced fractional excretion of electrolytes | 423±172 | (16) |
| Type-2 diabetes patients (5 mg/day) | Improved endothelial function measured by brachial flow-mediated dilation | 779±186 | (16) |
| Rat acute kidney injury model (1 to 3 mg/Kg)<sup>a</sup> | Reduced serum creatinine and urea<br>Reduced interstitial hemorrhage<br>Reduced albumin leakage into renal tissue<br>Improved renal PECAM-1 staining | 867±295 to 2600±884 | (15) |
| Pig coronary ischemia reperfusion (1 mg/kg) | Improved vascular reactivity of microcirculation (coronary flow reserve) | 2080±166 | (15) |
| Endothelial cells | Barrier integrity (impedance)<br>Phosphorylation of ERK and AKT | EC<sub>50</sub> = 592<br>EC<sub>50</sub> = 171 – 277 | (15) |

<sup>a</sup> 1 mg/kg concentrations calculated by linear extrapolation from 3 mg/kg

S1P_1_ is also expressed directly on the cardiomyocyte and S1P_1_ activation causes heart rate reduction (8). In principle, a reduction in heart rate could be associated with cardio-protective effects, especially as cardiac overexpression of S1P_1_ has been shown to improve outcomes after experimentally induced myocardial infarction or stroke (27). However, SAR247799 has a low volume of distribution in rats and humans and this property limits concentrations in the heart relative to the vasculature (15). Indeed, SAR247799 has endothelial-protective effects in type-2 diabetes patients under conditions where there were only small changes in heart rate (16). Similarly, in the current studies SAR247799 caused 4 and 6% reductions in heart rate in the young- and old ZSF1 rats, respectively. These levels of heart rate reduction are unlikely to explain the cardioprotective effects seen in the present work. In addition, heart rate-lowering drugs have not demonstrated benefit in patients with HFpEF (31), lending support for a more endothelial-mediated mechanism.

Chronic kidney disease is prevalent in HFpEF patients and is associated with increased morbidity and mortality (32). The co-existence of chronic kidney disease with HFpEF may be due to common risk factors such as diabetes, or to the interaction of the renal and cardiac systems with each other (i.e., heart failure inducing renal hypoperfusion and dysfunction, or, conversely, renal failure causing cardiac dysfunction via increased afterload) (32). Ob-ZSF1 rats inherently develop renal disease and this models the cardiorenal syndrome of HFpEF patients (33). In the present study, Ob-ZSF1 rats developed age-dependent polyuria and proteinuria. There was also an increase in the FENa and FECl, in old ZSF1 rats. Inappropriate urinary Na^+^ and Cl^-^ loss indicates renal salt wasting which is common in CKD and hypertensive patients (34), and SAR247799 tended to decrease such losses in Ob-ZSF1 rats. Such action of SAR247799 could be by restoring glomerular-tubular balance, i.e. reducing salt delivery to the proximal tubules and subsequent reabsorption through improving the functional capacity per nephron (34). Indeed, SAR247799 improves endothelial function at the glomerular level and reduces the severity of acute kidney injury in rats (15), suggesting that it could have acted by improving renal tubules and their re-absorptive mechanisms in the present study.

Aged, male, obese ZSF1 rats are considered a useful preclinical model for investigating potential therapeutics for LVH, diastolic dysfunction and thus HFpEF, because they combine the obesity/metabolic phenotype with hypertension and kidney disease, and thus mimic well the setting of the HFpEF patient beyond solely the diastolic dysfunction (20). Various pharmacological agents with potential in HFpEF have been tested in Ob-ZSF1 rats. HFpEF cardiomyopathy is reduced in Ob-ZSF1 rats by blood pressure lowering agents, such as a AT_1_/ETa receptor blocker (35). Others have pointed on the action of glucose lowering agents such as empagliflozin (36) to improve diastolic dysfunction and cardiac hypertrophy, respectively. However, clinical studies have reported detrimental or no effect of these approaches in HFpEF patients (37), suggesting that other options should be investigated. Importantly, SAR247799 improved diastolic function independent of any blood pressure modulation (35, 36), a feature that distinguishes it from previous molecules tested in Ob-ZSF1 rats and from renin-angiontensin-aldosterone pathway modulators that have failed to show benefit in HFpEF patients (38). SAR247799 displays here a unique profile of beneficial effects in young and old Ob-ZSF1 rats, without affecting blood pressure or obesity. SAR247799 is not a blood pressure lowering agent as it did not reduce hypertension in treated ZSF1 rats in the present study and it did not modify blood pressure in diabetic patients in a previous study (16). Similarly, SAR247799 cannot be classified as an anti-diabetic drug as in this study and in the previous study in ZDF rats and in diabetic patients (16), it did not lower hyperglycemia. SAR247799 is also the first agent to show diastolic function improvements in Ob-ZSF1 rats by echocardiographic assessment (E/e’). E/e’ is a reliable non-invasive diastolic index translatable to clinical practice compared to invasive methods including tau and left ventricular end-diastolic pressure, as used in other molecules evaluated in Ob-ZSF1 rats (39). E/e’ is generally assumed to be less sensitive to preload than other echocardiographic indices of diastolic dysfunction such as E/A and therefore yields more accurate estimations of filling pressures (40). Agents aiming to improve the physiology of endothelium in this complex microvascular pathology remains one of the most promising approach because of the capability of endothelial-driven molecular pathways to reduce myocardial fibrosis, stiffness and dysfunction (7). Importantly, this is the first time that S1P_1_ activation has been pharmacologically associated with benefits in a model of LVH and diastolic dysfunction. Thus, SAR247799 is a promising therapeutic approach and deserves to be tested in the clinical context of LVH and diastolic dysfunction, major components of HFpEF.

Nevertheless, the present study had several limitations. 1) Despite adopting the same dosing regimen, steady state plasma concentrations of SAR247799 were 7-fold lower in the older cohort compared to the younger cohort. We could speculate that age of the animals could have affected absorption, metabolism or excretion parameters. However, plasma exposure in older animals were still in a similar range to concentrations in our previous published study in ZDF rats where we demonstrated endothelial protection and were not associated with significant lymphocyte reduction, a marker of S1P_1_ desensitization (16). In both cohorts, heart rate reductions were minimal (4-6%) and there was no evidence for S1P_1_ desensitization (i.e. no significant lymphocyte reduction) following 4-week treatment. Although the more impressive effects of SAR247799 were achieved in the older cohort, it is still possible that sub-optimal concentrations could have produced sub-optimal efficacy. 2) Four-week treatment duration may not have been sufficient to see changes in microvessel density, and a longer duration study may have been warranted to see effects through this mechanism. 3) For ethical reasons, we did not study mortality in the rat study which is a key outcome in clinical trials in HFpEF. 4) We did not evaluate the cardiac phenotype and the effect of SAR247799 in female rats. This represents a limitation of the study as the prevalence of HFpEF is higher in women than in men. 5) E/A analysis by doppler echocardiography was not feasible in **old ZSF1 rats** because of the high heart rate that lead to E and A wave fusions. Speckle tracking echocardiography for the evaluation of left ventricle and atrial strain was not performed in this study. 6) Despite invasive in both animals and human, the hemodynamic evaluation of left ventricular end-diastolic pressure remains the gold standard of HFpEF and was not here assessed. 7) Finally, anti-hypertrophic effect and internal pathway of S1P_1_ activation through SAR247799 were not addressed in this study.

## 5. Acknowledgements

In vivo experiments were conducted by MFE, XC, SB and FG. Analyses were done by MFE (echocardiography, biological parameters) and XC (telemetry, SBP variability). PK determination was performed by AR, and histology by AML. MFE, BP, AAP, AC, MPPH and PJ conceived and designed the studies. MFE, BP and AAP wrote the manuscript. All authors approved the final version.

We would like to thank the *Translational In Vivo Model – In Vivo Research Support Services* of Sanofi R&D for animal caregiver and technical support. We thank Dorothée Tamarelle for support in statistical analysis. We are also thankful to Carole Legeay and Sylvie Bethegnies for pharmacokinetic measurements and Jean-Pierre Ripoll et Christophe Frances for plasma samples reception and storage. Finally, we would like to thank Christophe Gros and Naimi Souâd for histological analysis support and Cécile Canalis for contribution in data collection.

---

**Supplementary Figure** 1:

**(A)** E/A, **(B)** A wave at mitral valve. Parameters were measured in young animals. In old animal data were not determined (nd).

Data are expressed as mean ± SEM, †p<0.05, comparison of Le-ZSF1-CTRL to Ob-ZSF1-CTRL using a Student t-test. Le-ZSF1-CTRL N=9, Ob-ZSF1-CTRL N=11 and Ob-ZSF1-SAR247799 N=11

**Supplementary Figure 2**: Representative echocardiographic images of E wave, e’ wave and left ventricular thickness in (A), (B) and (C) for young animals and in (D), (E) and (F) for old animals, respectively.

